# Single and complex spikes relay distinct frequency-dependent circuit information in the hippocampus

**DOI:** 10.1101/2022.04.06.487256

**Authors:** Eric Lowet, Daniel J. Sheehan, Rebecca A. Mount, Sheng Xiao, Samuel L. Zhou, Hua-an Tseng, Howard Gritton, Sanaya Shroff, Krishnakanth Kondabolu, Cyrus Cheung, Jerome Mertz, Michael E. Hasselmo, Xue Han

## Abstract

Hippocampal neurons generate either single spikes or stereotyped bursts of spikes known as complex spikes. Although single and complex spikes co-occur in the same neuron, their contribution to information processing remains unclear. We analyzed hippocampal CA1 neurons in awake mice and in behaving rats, combining cellular membrane voltage imaging with optogenetics and extracellular recordings. We found that network-driven subthreshold membrane rhythms in the theta versus gamma frequencies preferably entrained complex versus single spikes in individual neurons. Optogenetic membrane perturbation revealed a causal link between subthreshold theta and gamma power and the initiation of complex versus single spikes. Further, single and complex spikes exhibited different place field properties and frequency-dependent coding during spatial navigation. Thus, individual hippocampal neurons do not integrate theta and gamma rhythms into a combined spike timing code, but instead, transmit frequency-specific information as distinct output modes of single versus complex spikes during spatial cognition.

## Main

Action potentials carry the information output of a neuron. Action potential initiation is determined by subthreshold membrane voltage (Vm) at the soma that reflects a spatiotemporal transformation of synaptic inputs shaped by intrinsic membrane ionic conductances^1,2^. Not only is the absolute Vm critical for action potential generation, but the temporal dynamics of Vm also impact the timing of action potentials, also known as spike timing^3^. Vm is shaped by coordinated circuit activity. One way to capture coordinated network activity is to record extracellular local field potentials (LFPs), in which synchronized synaptic inputs are reflected as LFP rhythms. In hippocampal CA1 circuits, LFPs have prominent theta^4–8^ and gamma rhythms^9–13^, especially during spatial navigation. CA1 receives two major synaptic input sources that originate from CA3 and the entorhinal cortex layer 3 (EC3). CA3 and EC3 inputs converge onto CA1 pyramidal neurons in a theta phase coordinated manner^10,11,14^. Additionally, these inputs exhibit different gamma rhythms across the slow (30-50Hz), middle (60-100Hz) and fast (>100Hz) frequencies and are thought to drive layer-dependent CA1 gamma rhytms^10,11^.

Extracellular recordings in the brain have identified one prominent spike timing feature, spike bursts (SB)^15^, meaning that several spikes occur within a short period or with short inter-spike-intervals (ISIs). SB has been broadly observed in many types of neurons across cortical and subcortical brain regions, and implicated in a variety of neural circuit functions, including signal transmission^16,17^, plasticity^18–21^, multiplex coding^3,22–25^ and integration of different input streams^26–28^.

Extracellularly recorded SB *in vivo* has often been related to intracellularly recorded complex spikes (CS) *in vitro*, where high density spikes occur during strong and long-lasting Vm depolarization. Thus far, a limited number of intracellular studies confirm that CS is associated with sustained and strong Vm depolarization in the brain of living animals^20,21,29–31^. Theoretical studies and *in vitro* brain slice experiments suggest that sustained Vm depolarization underlies CS generation, leading to some important questions. Such as, how do the sustained Vm depolarizations arise during behavior? Do they relate to specific input dynamics captured by the LFPs? Critically, compared to single spikes (SS), does CS represent a unique neural circuit output modality, and does CS carry behaviorally or contextually different information?

To understand how CS and SS are differentially regulated by subthreshold membrane voltage dynamics and network inputs, and whether CS represents a unique spiking output mode different from SS, we performed high-speed SomArchon voltage imaging from individual CA1 neurons, and compared intracellular voltage dynamics with simultaneously recorded LFPs in awake mice. To link CS and SB, we further compared our voltage imaging findings in mice to extracellular recordings obtained in rats during spatial navigation.

### High-speed voltage imaging of single spikes (SS) and complex spikes (CS) in individual CA1 neurons

To perform single cell voltage imaging and optogenetics in dorsal hippocampal CA1, we surgically implanted an imaging window coupled with an infusion cannula and an LFP electrode, and infused AAV9-Syn-SomArchon-p2A-CoChR-Kv2.1 through the infusion cannula into CA1. During recording, habituated animals were head-fixed under the microscope objective on a floating Styrofoam ball supported by air pressure (**Fig. 1a**). Voltage imaging of SomArchon expressing neurons was obtained with a high-speed sCMOS camera using a 40x water immersion objective. To examine how subthreshold membrane potential (Vm) influences spike timing, we removed identified spikes from SomArchon traces to obtain the Vm trace for each neuron (see Methods). We observed that many neurons exhibited both intermittent SS and bursts of spikes that are associated with substantial Vm depolarization corresponding to events known as CS. To better characterize the action potential features of SS and CS, we performed ultra-fast voltage imaging at 5-10kHz (**Fig. 1b**). We quantified the full-width at half-maximum-amplitude (FWHM) of each action potential, and found that the first action potential within a CS had a similar FWHM as that of SS. However, the FWHM gradually widened for the subsequent action potentials within a CS (**Fig. 1c-d**), increasing from 0.95ms (1^st^ spike) to 2ms (5^th^ spike or more). This increase in FWHM was accompanied with a decrease in spike amplitude (**Fig. 1e**), similar to prior observations, and thus confirmed that SomArchon voltage imaging has sufficient sensitivity to capture individual CS^20^.

**Fig.1:**
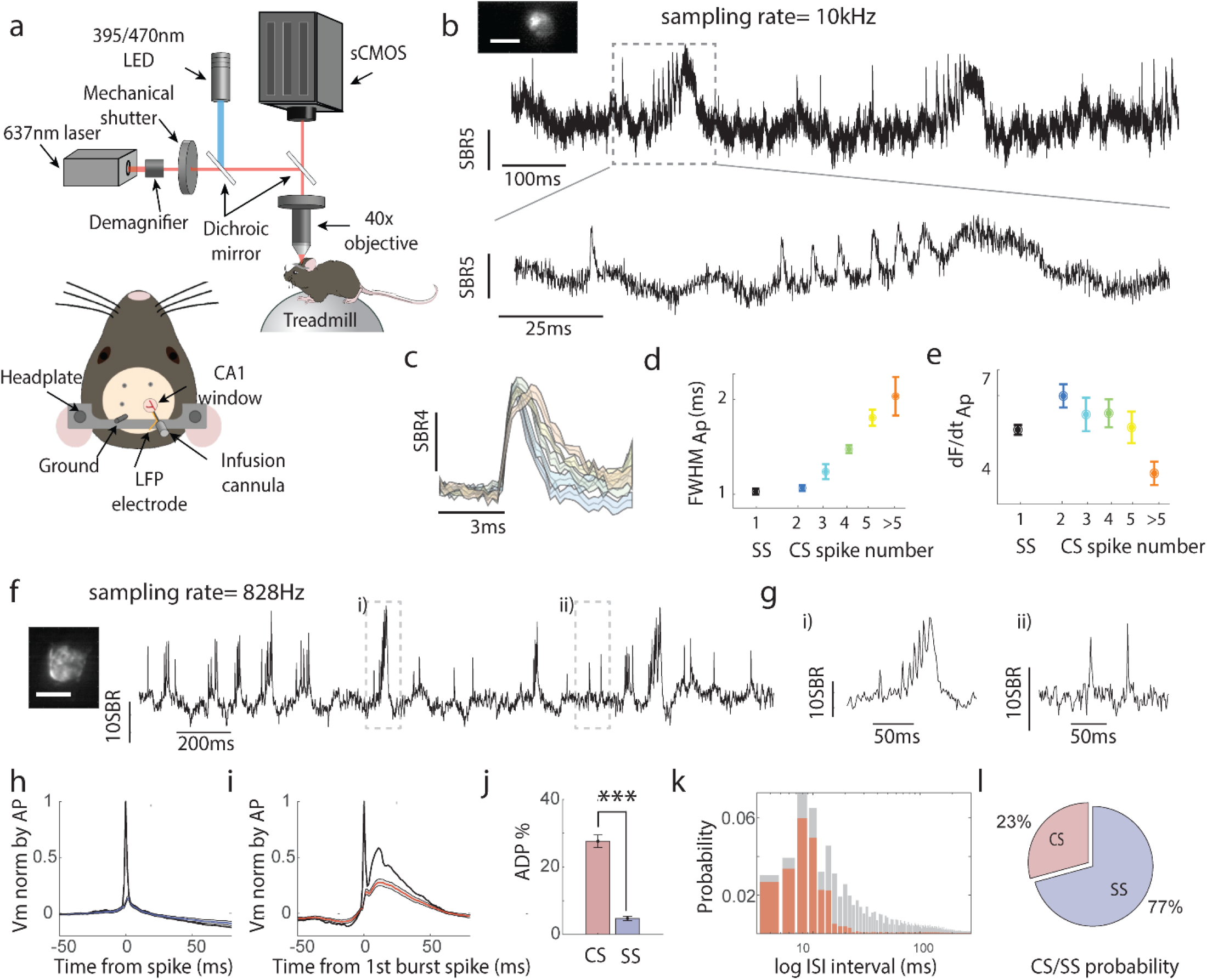
High-speed voltage imaging characterization of CA1 single spikes (SS) and complex spikes (CS) in awake head fixed mice. **(a)**. Illustration of SomArchon voltage imaging and animal preparation. **(b)**. An example SomArchon fluorescence recording at 10kHz from a CA1 neuron. Top, average SomArchon fluorescence at spike peak for the recorded neuron. Scale bar, 25μm. Bottom, SomArchon fluorescence trace, and Zoom-in view of the period highlighted by the dashed box. **(c)**. SomArchon fluorescence aligned to the peak of APs within CS recorded at 5-10KHz. The colors correspond to the sequence of APs, with blue being the 1^st^ AP; light blue, 2^nd^; green 3^rd^; yellow 4^th^; orange 5^th^ or more. **(d)**. Quantification of full width at half maximum (FWHM) for APs within SS and CS. FWHM increased as a function of CS spike number (linear regression slope, p=1e^-20^, n=101 spikes). **(e)**. Quantification of spike amplitude calculated as maximum SomArchon fluorescence change (dF/dt) around each AP within SS and CS. Spike amplitude decreased as a function of CS spike number (linear regression slope, p=0.0056, n=101). **(f)**. An example 3-second-long SomArchon recording at 828Hz from a CA1 neuron. Left, average SomArchon fluorescence at spike peak for the neuron recorded. Scale bar, 15μm. Dashed lines represent zoom-ins shown in g. **(g)**. Zoom-in of the example trace segments indicated in f. (**h-i**). Spike-triggered SomArchon traces, either triggered to the peak of single spikes (SS) shown in **(h)** or to the peak of the 1^st^ spikes of CS **(i)**. The black line corresponds the spike-triggered averaged raw SomArchon fluorescence trace. The blue or red line with the shaded area corresponds to SomArchon trace with spikes removed (Vm). **(j)**. Quantification Vm after-spike depolarization potential (ADP), after SS or CS (5ms-25ms after spike peak). Shown are ADP normalized to pre-spike period (25ms to 5ms before spike peak). ADP was stronger in CS than SS (paired t-test, p<1e-20, df=66). **(k)**. The inter spike interval (ISI) distribution of all spikes (gray) including CS (red). **(l)**. Overall percentage of spikes classified as CS or SS (n=67 neurons).

After confirming the basic features of action potentials within CS and SS using ultrafast 5-10kHz imaging, we imaged many more neurons at a lower speed of 828Hz for enhanced SomArchon signal quality^32^ (**Fig.1f-g, Extended data Fig.1**) while capturing Vm depolarizations. Compared to a SS, the first spike within a CS was followed by a strong Vm after-spike depolarization (ADP), and the entire CS duration was accompanied with substantial Vm depolarization (**Fig.1h-i**). To separate SS and CS for subsequent analysis, we thus employed both Vm ADP and inter spike intervals (ISI) as criteria (**Fig.1j-k**). *In vivo* extracellular recording studies typically used ISI thresholds of 6-10ms and recent intracellular studies have shown that

ISIs between spikes within a CS can be considerably longer^20^, and thus we used a combination of ISI <14ms and an ADP of >15% (4SD of SS distribution) as thresholds to classify each spike as CS versus SS. Using these thresholds, we found that around 23% of all recorded spikes were CS (**Fig.1l**), and 82% of neurons had at least 10% CS spike probability (49% of neurons with >20% CS probability and 13% with >40%CS probability).

### CS and SS exhibit different relationships to Vm oscillations at theta versus gamma frequencies

The long-lasting ADP associated with CS occurred on a time scale of multiple tens of milliseconds (**Fig. 1I**), notably corresponding to the theta oscillation time scale. Thus, we next explored the relationship between CS ADP and Vm wavelet spectral power (**Fig.2a**). We found that CS was associated with a strong increase in Vm power at theta frequencies (3-12Hz), whereas SS was weakly related to Vm theta power. Over the neural population, we detected significantly stronger Vm theta power around CS than around SS (**Fig.2b**). The selective association of CS with Vm theta oscillations was further confirmed using time-domain analysis (**Extended data Fig.2a**), where we aligned theta frequency filtered Vm to the 1^st^ CS spike or SS. These results demonstrate that CS is selectively associated with increased Vm theta power, due to the prominent theta-frequency time scale Vm depolarizations that are specific to CS.

**Fig.2:**
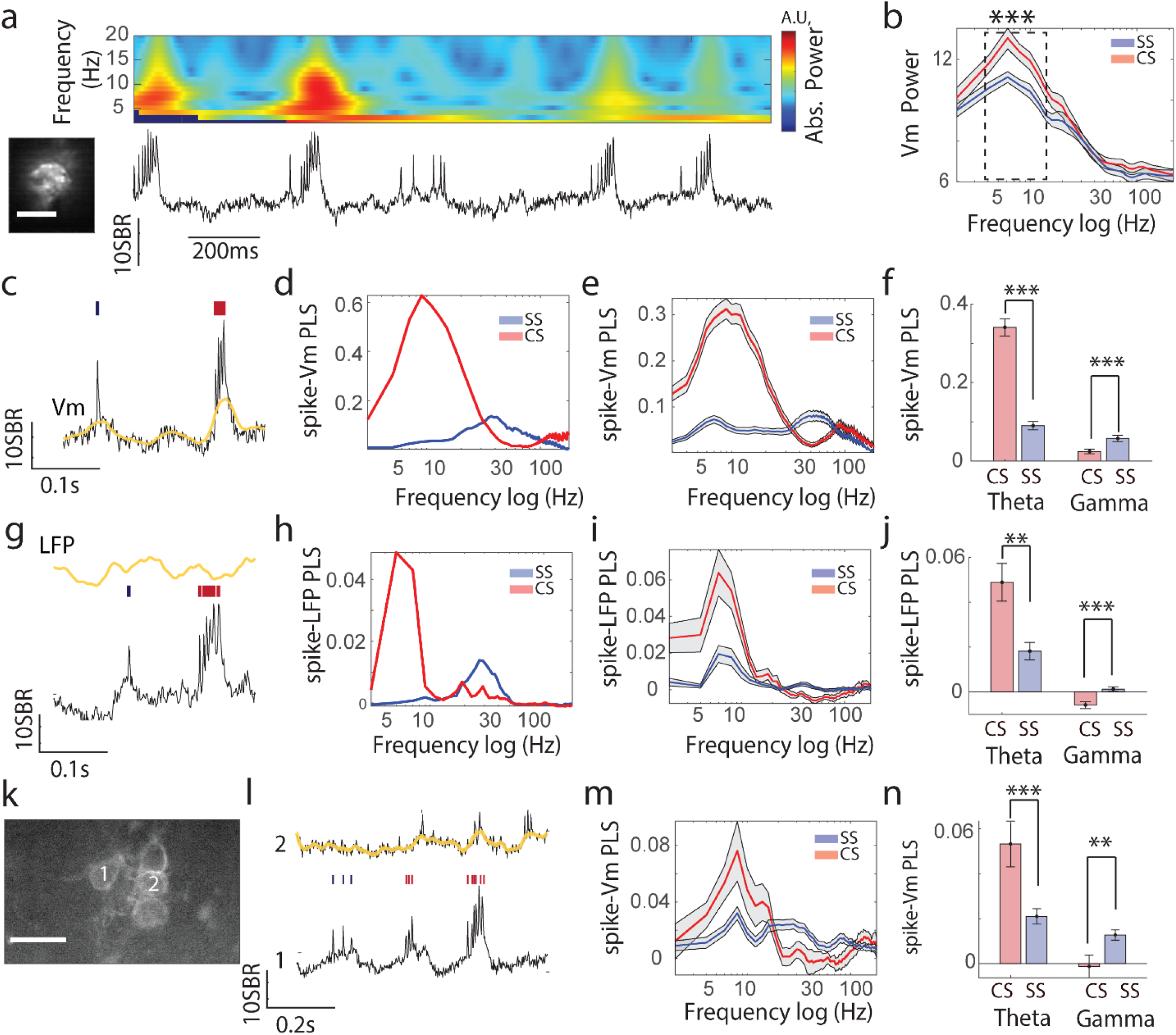
SS and CS exhibit distinct relationship to Vm and LFP oscillations at theta and gamma frequencies. **(a)**. An example SomArchon trace of a CA1 neuron with the corresponding Vm wavelet power spectrum on top. Left, Average SomArchon fluorescence intensity at spike peak in the neuron recorded. Scale bar, 15μm. **(b)**. Vm spectral power aligned to SS (blue) and the 1^st^ spikes within CS (red). Vm theta power is greater for CS than SS (paired t-test, p= 9.13e^-8^, df=43). **(c-f)**. Spike timing relative to Vm phase for SS and CS. **(c)**. Example SomArchon trace of a CA1 neuron with both SS (blue) and CS (red). Yellow is the theta-frequency filtered Vm trace. **(d)**. Phase-locking strength (PLS) of spikes to Vm across frequencies for CS (red) and SS (blue) for an example neuron. **(e)**. Mean PLS of spikes to Vm across frequencies for the population of recorded neurons (n=49). **(f)**. PLS of spikes to Vm theta (3-12Hz) and gamma oscillations (30-90Hz) for CS (red) and SS (blue). Spike-Vm theta PLS for CS is greater than for SS (paired t-test, p=1e^-20^, df=48). Spike-Vm gamma PLS for SS is greater than CS (paired t-test, p=1.4e^-7^, df=48). **(g-j)**. same as **c-**f, but for spike timing relative to simultaneously recorded LFPs. **(j)**. Spike-LFP theta PLS for CS is greater than for SS (paired t-test, p=0.0047, df=48). Spike-LFP gamma PLS for SS is greater than CS (paired t-test, p=8.33e^-5^, df=48) **(k-n)**. Multi-neuron SomArchon recordings using patterned illumination voltage imaging. **(k)**. Example field of view showing GFP fluorescence from 5 simultaneously recorded neurons expressing SomArchon-GFP. Scale bar 30μm **(l)**. Example SomArchon traces from a pair of neurons 1 and 2 shown in **k**. Spike-Vm phase coordination between CA1 neurons was considered between SS (blue) or CS (red) of one neuron and Vm (yellow trace) of the other neuron **(m)**. Similar as in **e**, PLS of SS or CS spikes from one neuron to Vm of another simultaneously recorded neuron, instead of Vm of the same neuron as in **e**. (**n)**. Spike-Vm PLS in theta range (3-12Hz) and gamma range (30-90Hz). CS spikes show stronger PLS to other neurons’ Vm theta oscillations than SS (independent t-test, p <1.97e^-4^, df=48, only CS and SS with >10spikes were included). SS in contrast exhibits stronger PLS to another neuron’s Vm gamma oscillations (independent t-test, p= 0.006, df=79).

We then investigated how Vm theta oscillations organize the timing of spikes within SS and CS. We calculated spike-Vm phase-locking strength (PLS, see Methods), a measure of temporal consistency of spike occurrence at a specific phase of the Vm oscillation adjusted for spike number. Specifically, we compared the PLS of each individual spike within SS or CS relative to Vm theta oscillation phase in each neuron, and then averaged across all neurons **(Fig.2c)**. We observed that CS exhibits significantly greater spike-Vm PLS at theta frequencies than SS, demonstrating that spikes within CS occur at a more consistent phase of the theta cycle than SS (**Fig.2d-f**). This result was also observed when considering the first CS spikes only (**Extended data Fig.3)**. Further, CS with more spikes showed stronger spike-Vm PLS at theta frequencies (**Extended data Fig.4)**. Thus, there is a preferential association of CS spikes with theta oscillations, and that the strength of phase locking depends on the number of spikes within a CS event.

Interestingly, even though SS exhibited relatively weak PLSs to Vm at theta frequencies, SS showed strong PLSs at gamma frequencies (30-90 Hz) (**Fig.2d-f**). Spike-Vm PLS at gamma frequencies^33^ is also a common feature of the hippocampus. At gamma frequencies, SS showed significantly stronger PLS than CS. Spike-Vm PLS at gamma frequencies calculated with only the first CS spike was also weaker than that of SS (**Extended data Fig.3**). We further noted that CS exhibited stronger PLS in the high-gamma frequency range (100-150Hz), which likely arose from the short ISI between spikes within CS. Together, at the single cell level, CS exhibited a stronger phase relationship to Vm theta and high-gamma oscillations, whereas SS exhibited a more consistent phase relationship to gamma rhythms.

### CS and SS are temporally coordinated at the network level

Theta and gamma rhythms are commonly observed in hippocampal LFPs and are linked to a variety of behaviors including working memory, novelty exploration, and memory recall. LFPs measure net extracellular transmembrane currents at the electrode recording site^34^, and LFP oscillations reflect coordinated currents. Because of the brief nature of action potentials, it has been suggested that synchronized slower synaptic currents are the main source for LFPs. Given that subthreshold Vm reflects the summation of soma-dendritic synaptic inputs, the distinct CS and SS relationship with Vm oscillations might reflect differential coordination with synchronized network inputs at different frequencies. We therefore examined how the timing of spikes within SS and CS in individual CA1 neurons relate to the phase of simultaneously recorded hippocampal LFPs (**Fig.2g**). We found that CS had stronger spike-LFP PLS at theta frequencies compared to SS (**Fig.2h-j**), and PLS was stronger for CS containing a greater number of spikes (**Extended data Fig.4**), similar to that observed with spike-Vm theta phase-locking. In contrast, SS exhibited stronger spike-LFP PLS at gamma frequencies, also consistent with that observed with spike-Vm gamma phase-locking (**Fig.2h-j**).

The distinct properties of CS and SS phase-locking to LFP theta versus gamma oscillations suggest that the two spike output modes might arise from differential coordination between the neuron and network inputs. Monopolar LFP, as used here, presumably reflects both local and distant current sources within a large tissue volume, and thus it is unclear whether it explicitly captures coordination within local CA1 neural assemblies. To examine how Vm of individual neurons relate to other nearby neurons, we imaged multiple CA1 neurons simultaneously using a patterned illumination system for larger-scale voltage imaging ^32^. To avoid Vm fluorescence signal cross-contamination, we only analyzed neuron pairs separated by at least 40 microns, where Vm cross-contamination between simultaneously recorded neurons is negligible^32^. We aligned theta filtered (3-12Hz) Vm of a given neuron to SS or CS of nearby neurons and computed spike-Vm PLS between neurons (**Fig.2k**). Across neuron pairs, we found that CS from one neuron exhibited stronger PLS with Vm theta oscillations of another neuron than SS, whereas SS exhibited stronger PLS in the gamma range than CS (**Fig.2l-n**). CS and SS are thus preferentially linked to distinct subthreshold membrane voltage dynamics at the single cell level measured by Vm, and to different network oscillations measured by LFPs. These observations further confirm that CS and SS are unique output patterns that are differentially coordinated by different rhythms at the network level.

### Distinct Vm dynamics causally determine CS versus SS spiking output patterns

The distinct relationship of SS and CS to Vm and LFP theta versus gamma oscillations suggests that the two spiking modes are correlated with distinct input patterns. Since action potential initiation is dictated by Vm dynamics, we next directly tested the causal relationship between membrane input dynamics and the probability of CS or SS occurrence. We performed SomArchon voltage imaging while optogenetically depolarizing neurons that co-expressed CoChR either using continuous or rhythmic optogenetic stimulation.

We first tested how non-rhythmic membrane depolarization alters Vm oscillations and spike modes using continuous 1.5-second-long blue light to activate CoChR (**Fig.3a, Extended data Fig.5**). Upon stimulation, we detected a transient increase in the population spike rates of both SS and CS (**Fig.3c&d**) that lasted for ∼200ms (**Fig.3e**), which was followed by a sustained (0.2-1.5s) increase in SS firing rate, but not CS firing rate (**Fig.3f**). In line with the prominent increase in SS firing rate, population Vm power increased in the wider gamma frequency range (>35Hz) with a peak centered at ∼50-55Hz (**Fig. 3g**). Similarly, because CS firing rate did not consistently increase as a population, Vm power at theta frequencies (3-12Hz) remained constant (**Fig. 3g**). However, we observed considerable variability in optogenetically induced CS firing rate change across individual neurons, with some neurons having increased CS and others decreased CS. We therefore further explored the relationship between changes in Vm power and CS spike rate at the individual neuron level. We found that optogenetically induced changes in CS firing rate correlated with changes in Vm theta power (3-12Hz, **Fig.3h-i**), whereas optogenetically induced changes in SS firing rate correlated with changes in Vm gamma power (30-90Hz, **Fig.3h-i**). Together, dynamic switching between CS and SS firing modes closely reflects the underlying Vm frequency content.

**Fig.3:**
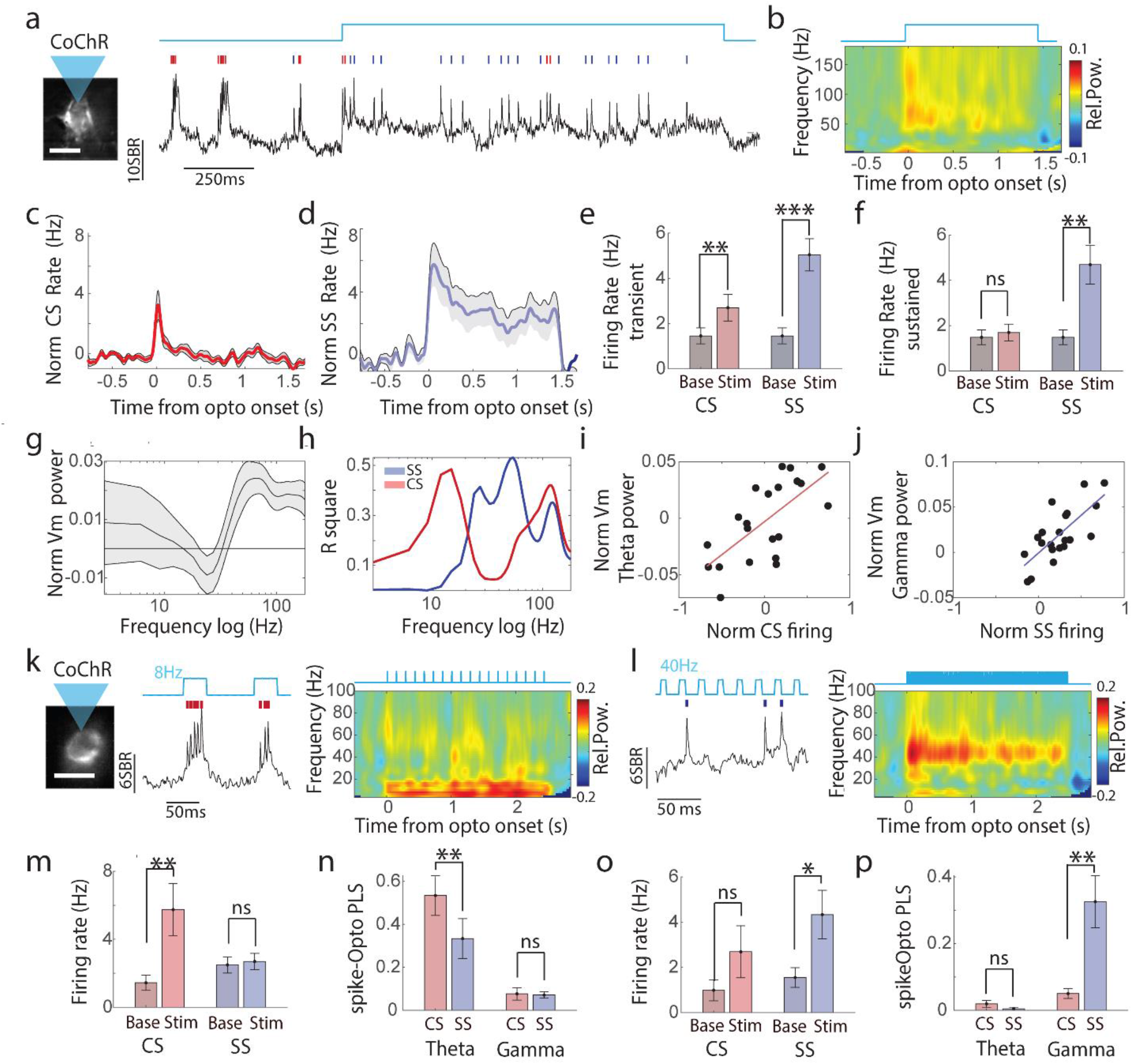
SS and CS respond differently to varying optogenetic stimulation patterns. **(a)**. An example SomArchon trace from a CA1 neuron expressing both SomArchon and CoChR, before, during and after 1.5-second-long continuous focal blue light illumination. CS and SS are marked with red and blue lines respectively. **(b)**. Population trial-averaged (n=25) wavelet power spectrogram of Vm. Vm Power is normalized to the 0.5 second baseline period before optogenetic stimulation onset. **(c, d)**. Population CS (**c**) and SS (**d**) firing rate probability upon continuous optogenetic stimulation. **(e)**. Transient change in population firing rate for CS and SS within 200ms of optogenetic stimulation onset. Both CS (paired t-test, p=0.007, df=23) and SS (paired t-test, p=1.9e^-4^, df=24) showed an increased in firing rate relative to the 0.5 second baseline period before stimulation onset. **(f)**. Sustained change in population firing rate for CS and SS during the 0.2-1.5 second time window after stimulation onset. SS (paired t-test, p=0.003, df=24), but not CS (paired t-test, p=0.96, df=23), showed a significant increase in firing rate. **(g)**. Optogenetic stimulation induced population Vm power change relative to 0.5-second long baseline before stimulation onset ((Vm(stimulation)-Vm(baseline))/(Vm(stimulation)+Vm(baseline)), n=26 neurons). **(h)**. The coefficient of determination (R square) obtained from linear regression between optogenetic-evoked spike rate change of CS (red) or SS (blue) for a given neuron and the optogenetic-evoked change of Vm power across frequencies. **(i)**. Scatter plot of optogenetic-evoked CS firing rate change and Vm theta power change (5-12Hz) per neuron. There was a significant positive correlation (linear regression, r^2^=0.45, p=0.0009, n= 22). When using SS firing rate, the relationship was not significant (linear regression, r^2^= 0.02, p=0.57, n=22). **j**, Scatter plot of optogenetic-evoked SS firing rate change and Vm gamma power change (30-70Hz). There was a significant positive correlation (linear regression, r^2^=0.49, p=0.0003, n=22). When using CS firing rate, the relationship was not significant (linear regression, r^2^=0.12, p=0.13, n=22). **(k, l)** SomArchon trace from an example neuron upon 8Hz (**k**) and 40Hz (**l**) pulsed optogenetic stimulation. CS and SS are marked with red and blue lines respectively. **(m)**. Quantification of firing rate for CS (red) and SS (blue) before (base) and after each stimulation light pulse (stim) delivered at 8Hz. **(n)**. Quantification of spike PLS to stimulation pulse patterns at 8Hz for CS (red) and SS (blue). PLS for CS is significantly higher than for SS (paired t-test, p=0.0091, df=10). **(o)**. Same as **m** but for light pulse delivered at 40Hz. Firing rate for SS is significantly higher after each stimulation pulse compared to the baseline before each light pulse (paired t – test, p=0.027, df=10). **(p)**. Quantification of Spike-Vm PLS to stimulation pulse pattern at 40Hz. PLS for SS is significantly stronger than for CS (paired t-test, SS, p=0.0056, df=7).

Next, to directly evaluate how rhythmic Vm depolarization influences CS and SS, we pulsed blue light at 8Hz (**Fig.3k)** or 40Hz (**Fig.3l**). During 8Hz stimulation, only the firing rate of CS increased (**Fig.3m**), whereas during 40Hz stimulation, only the firing rate of SS increased (**Fig. 3o**). To further quantify the timing of CS and SS relative to stimulation pulses, we calculated the PLS of SS and CS to blue light stimulation pulses. We found that CS (all spikes considered) had a stronger spike PLS to 8Hz stimulation pulses than SS (**Fig.3N**), whereas SS had a significantly greater PLS to 40Hz stimulation pulses than CS (all spikes considered) (**Fig. 3p**). Thus, optogenetic enhancement of Vm theta oscillations preferentially increased the probability of CS, and CS occur at a defined phase of Vm theta oscillations. In contrast, optogenetic enhancement of Vm gamma oscillations resulted in preferential spiking in SS mode, and SS occurrence was temporally aligned to stimulation pulses. These findings provide direct evidence that Vm dynamics determine SS and CS occurrence, and organize spike timing of SS and CS. The fact that the same neuron can dynamically switch between SS and CS according to rhythmic membrane currents further confirms that SS and CS are distinct output modes influenced by input rhythmicity features.

### CA1 spike bursts (SB) and single spikes (SS) are differentially associated with LFP theta and gamma oscillations during spatial navigation in rats

Hippocampal CA1 is critical for spatial navigation and spatial memory^35,36^. Many CA1 neurons have place fields, specific regions of an environement where they spike, and CA1 spiking has been widely related to theta and gamma network rhythms that are dynamically modulated during learning and memory^35,37^. Our voltage imaging and optogenetic results demonstrated that many CA1 neurons have intermingled CS and SS, and the dynamic switching between CS and SS depends on Vm and LFP rhythms. These results predict that CS and SS might code different spatial information by selectively coordinating theta versus gamma network rhythms during navigation behavior. To test this hypothesis, we performed extracellular tetrode recordings in the dorsal CA1 of rats trained to run bidirectionally, inbound and outbound relative to a home box, on a custom-built wooden linear track in a room with environmental cues (**Fig.4a-b**). Since there is no Vm available from extracellular recordings, we separated spike bursts (SB) from single spikes (SS) using a traditional ISI-based criterium of <10ms alone (**Fig.4c**). We called them SB instead of CS, to distinguish the difference in recording modalities. Using this threshold, we found that around 30.5% of all recorded spikes were SB.

Similar to our intracellular voltage imaging results in mice, we observed that extracellularly recorded SB across the rat CA1 population exhibited several-fold stronger spike-LFP PLS at theta frequencies (5-12Hz) than SS (**Fig.4e-g**). Further, spike-LFP theta PLS increased for SB containing more spikes (**Extended data fig. 4**). In contrast, SS exhibited stronger spike-LFP PLS at gamma frequencies than SB (**Fig.4 e-g**), with a peak in the mid-gamma range (∼70-100Hz) that has been associated with EC3 inputs^11^. SB thus exhibited preferential phase consistency with LFP theta oscillations, whereas SS were more strongly associated with LFP gamma phase.

Our optogenetic studies demonstrated that rhythmic Vm inputs at theta versus gamma frequencies differentially promoted CS versus SS spike rate. Since LFP measures network dynamics that capture synchronized inputs, we reasoned that SB and SS between neurons would be coordinated by theta versus gamma oscillations respectively. To test this, we computed pairwise cross-correlograms between spike trains containing only SB or only SS from simultaneously recorded neurons. We found that SB spike train cross-correlograms had a stronger theta component (**Extended data Fig.6**), whereas SS spike train cross-correlograms had a stronger gamma component (**Extended data Fig.6**). These results highlight that SB and SS spiking output modes are temporally coordinated by distinct network rhythms within local cell assemblies, in line with our multi-neuron SomArchon voltage imaging results (**Fig. 2m, n**).

### SB and SS represent distinct behavioral codes during spatial navigation

CA1 theta and gamma oscillations have been linked to distinct generation mechanisms^4,13^, and thus these oscillations may carry different behavioral information as they are processed across hippocampal structures. The selective association of theta oscillations with SB and gamma oscillations with SS during spatial navigation predict that SB and SS would be differentially modulated by different input sources and thus exhibit distinct spatial coding properties. To explore this, we first compared how SB and SS firing rates are modulated by spatial locations by calculating the ratio of peak firing rate to mean firing rate for each neuron (**Fig. 4h-i**). Because place field structures of CA1 neurons vary depending on whether the rats ran in the inbound or the outbound direction^7^, we used the firing rate of each place cell when rats ran in the direction that exhibited the strongest modulation of spike rate calculated using all spikes (see Methods). We found that SB firing rate modulation was significantly greater than SS as animals ran through the place fields (**Fig. 4i**). To further quantify the spatial information coding ability of SB and SS, we computed the information theoretic measure to estimate the amount of information a spike contains about the animal’s position expressed as bits per spike (see Methods). We found that SB contained greater spatial information than SS (**Fig.4j**). Thus, not only are SB and SS firing rates differentially modulated by spatial locations, but these two spiking modes have varying spatial information coding ability.

**Fig.4:**
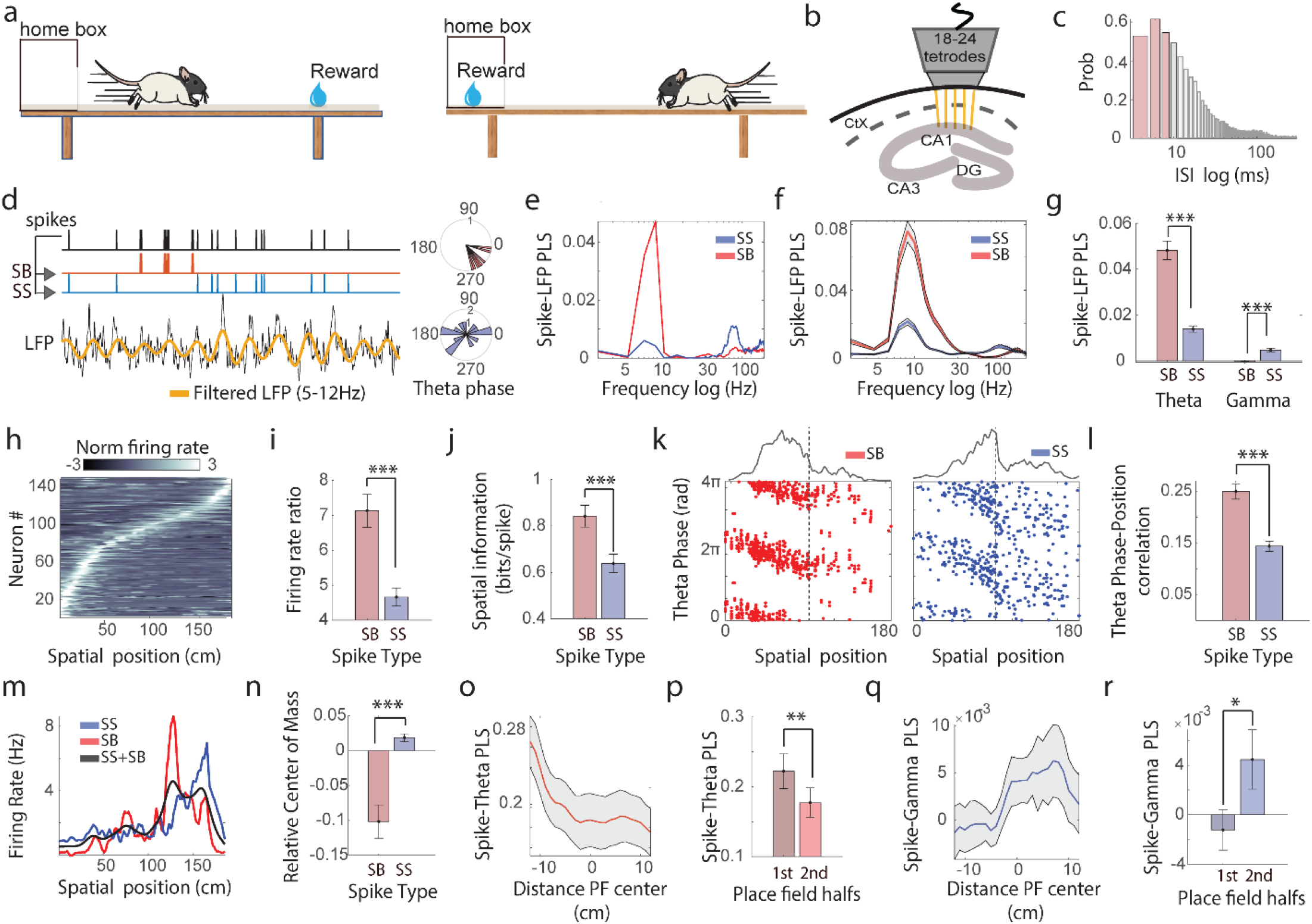
Distinct spatial coding and phase precession properties of CA1 single spikes (SS) and spike bursts (SB) during spatial navigation in rat. **(a)**. Schematic of the behavioral setup, in which rats were trained to run through a linear track in an outbound (left) and inbound (right) direction from the home box. **(b)**. Schematic of the extracellular tetrodes placed in the dorsal CA1. **(c)**. Inter-spike interval (ISI) distribution histogram across all spikes recorded in all CA1 neurons. ISIs correspond to spikes within single spikes (SS) versus spike bursts (SB) are depicted as gray and red respectively. (**d-g)**. Analysis of spike phase relative to LFP oscillations. **(d)** Left, an example LFP trace and simultaneously recorded spike train from an individual place cell. Every spike of the spike train (black) is classified as either SB (red) or SS (blue). The theta filtered LFP trace (5-12Hz) is in gold. Right, spike phase locking strength (PLS) to LFP theta oscillations plotted as polar histograms for SB (top) and SS (bottom). **(e)**. PLS of SS (blue) and SB (red) to LFP oscillations across frequencies for an example neuron. **(f)**. Population averaged PLS of spikes to LFP oscillations across frequencies (n=147 neurons) **(g)**. Quantification of PLS for SS and SB relative to LFP theta (5-12Hz) and gamma (30-100Hz) oscillations. SB PLS to LFP theta oscillations is significantly higher than SS (paired t-test, p<1e^-20^, df=146). SS PLS to LFP gamma oscillations is significantly higher than SB (paired t-test, p = 1.3e^-15^, df=146). **(h)**. Normalized z-scored firing rate (all spikes) of individual place cells as a function of animal position on the linear track. Neurons are sorted based on the timing of their peak firing rate respectively. **(i)**. The ratio of the peak firing rate relative to the mean firing rate for SB and SS. SB firing rate modulation was significantly stronger than SS (paired t-test, p=3.55e^-8^, df=146) **(j)**. SB and SS spatial information metric, calculated as bits per spike. (**k)**. An example CA1 neuron’s spike timing relative to LFP theta oscillation phase over animal’s spatial location for SB (red) and SS (blue). Each dot of the raster plot indicates a spike. **(l)**. Circular-linear correlation values between spike phase relative to LFP theta oscillations (5-12Hz) and animal’s spatial location. SB shows stronger correlations with spatial location than SS (paired t-test, p=1.77e^-13^, df=130). **(m)**. Example neuron’s spike rate plotted over spatial location, calculated using SB alone (red), SS alone (blue) or all spikes (black). **(n)**. The difference between neurons’ place field centers calculated using SB and all spikes (red), or between SS and all spikes (blue). The relative place field center of mass between SB and SS was significantly different (paired t-test, p=3.25e^-5^, df=146). **(o)**. population averaged spike PLS to LFP theta oscillations aligned to the SB place field center of each neuron (PF). **(p)**. Mean spike-LFP theta PLS during the 1st half of the place field (PF) as animals approach the PF center, and the 2^nd^ halve of the PF as animals leave the PF center. Spike-LFP theta PLS is significantly higher for the 1^st^ halve of the PF than the 2^nd^ halve of the PF (paired t-test, p=0.007, df=62). **(q-r)**. Same as **o** and **p**, but for spike PLS to LFP oscillations in the gamma range). Spike-LFP gamma PLS is significantly lower for the 1^st^ halve of the PF than the 2^nd^ halve of the PF (paired t-test, p=0.0396, df=62).

Another prominent phenomenon of CA1 spatial coding is that individual CA1 neuron’s spiking^7^ and Vm theta oscillations ^29^ show a systematic shift to earlier phases of LFP theta oscillations when animals traverse the place field of the neuron, a phenomenon known as phase precession^4,8^. Because our voltage imaging data showed that CS had stronger PLS to Vm and LFP theta oscillations than SS (**Fig. 2f, j**), we hypothesized that SB spikes would have stronger phase precession than SS. To directly examine this prediction, we calculated the preferred phase of SB (all SB spikes) and SS relative to LFP theta oscillations at each spatial location (**Fig.4k**), and computed the circular-linear correlation between the preferred spike-LFP theta phase and animal’s spatial position. We found that the SB spike-LFP theta phase exhibited much stronger correlations with spatial location than SS (**Fig.4l**), suggesting that theta phase precession coding is more strongly driven by SB.

Finally, to directly investigate the difference in SB and SS spatial coding properties, we separately calculated the firing rate of SB and SS at different locations. We found that SB often spiked earlier than SS for a given direction (**Fig.4m**). Direct comparison of this spatial difference between SB and SS firing rate using center of mass calculation revealed that SB firing was significantly earlier than SS by 22.3±8.7cm (mean ± standard error, **Fig.4n**), suggesting that SB were prevalent when an animal first entered the place field whereas SS were more prevalent when an animal was leaving the place field. Given SB and SS are preferentially linked to LFP theta and gamma oscillations respectively, we further examined whether spike-LFP phase locking exhibits similar spatial difference. Indeed, we found that SB-LFP PLS at theta frequencies was stronger in the first half of the place field (**Fig.4o&p)**, whereas SS-LFP PLS at gamma frequencies was stronger in the second half of the place field (**Fig.4q&r**). Thus, SB and SS contribute distinctively to neurons’ place field structures depending on network oscillation states. Together, these results demonstrate that SB, presumably corresponding to intracellularly recorded CS, and SS code different spatial information by selectively transmitting theta versus gamma network rhythms during navigation behavior, and SB plays a dominant role in theta phase precession.

## Discussion

Complex spikes (CS), and spike bursts (SB) more generally, have been widely documented in the hippocampus ^1–5^, various subcortical nuclei ^6–8^, the cerebellum ^9–11^ and neocortical circuits ^12–15^ across various species. Yet the role for CS in circuit processing and behavior remain largely elusive. Here, we studied the subthreshold membrane potentials (Vm) and spike output patterns from individual CA1 neurons in behaving mice using high-resolution SomArchon based cellular voltage imaging. We found that CS and SS co-exist in the majority of neurons analyzed, presumably pyramidal cells, occurring intermittently and at variable rates. These results provide direct experimental support that CS and SS are a common feature of CA1 neurons, consistent with previous *in vivo* CA1 patch clamp studies^16–19^ that have captured CS in a limited number of neurons in behaving animals.

The biophysical mechanisms for CS generation are not fully understood, but likely involve activation of specific voltage-dependent membrane currents^20^ and dendritic plateau potentials ^21–23^. In addition, CS and SS might critically rely on the timing of inhibitory synaptic currents that originate from an interplay between interneurons that target the soma versus the dendrites^23,24^. It is highly plausible that rhythmic inputs at theta versus gamma frequencies selectively recruit ionic and synaptic currents leading to these two distinct spike output modes.

Theta and gamma oscillations are prominent in the hippocampus during active behavior and are proposed to underlie network functions supporting memory and navigation^25–27^. Although LFP theta and gamma oscillations having been shown to be cross-frequency coupled^28–31^, our results demonstrate that theta and gamma oscillations are preferentially related to distinct spike outputs as CS and SS respectively. It therefore challenges the notion that theta and gamma form a dual oscillator code^29^, in which spike timing is precisely shaped by theta and gamma rhythms simultaneously. Instead, our data supports the notion that theta- and gamma-mediated network inputs to individual neurons dynamically switch between these two spike output modes. However, although SS exhibited a much less precise phase relationship with theta oscillation than CS, our data showed that SS timing remained modulated by theta oscillations.

CA1 neurons receive perisomatically targeted inputs from CA3 and apical dendritically targeted inputs from entorhinal cortex layer 3 (EC3). Theta rhythms are synchronized across different structures of the hippocampus, and CA3 and EC3 pathways activate CA1 neurons in a theta phase dependent manner^21,25^. Coordinated dendritic and somatic activation of CA1 neurons has been found to support CS generation^21,23,32^ and might explain why CS are strongly theta-locked. In contrast, gamma rhythms have been found to be locally generated and occurring at variable frequencies across CA1 layers and throughout different structures of the hippocampus^28^. CA3 and EC3 pathways to CA1 have been associated with slow (30-50Hz) and middle (60-100Hz) gamma frequencies respectively^28,33^. Our observations show that gamma frequencies preferentially entrain SS, suggesting that pathway-specific gamma-band communication^34–36^ is primarily carried by SS. Since CS is associated with strong dendritic and somatic membrane depolarization that leads to the ‘ballistic’ generation of high-frequency spikes (often >100Hz), the occurrence of CS likely prevents reliable entrainment of CS spike timing by gamma-rhythmic inputs.

Theoretical and experimental studies have shown that CS is associated with enhanced and reliable transmission to downstream targets^7,37,38^. SS, however, has been assumed to be less effective in inducing reliable postsynaptic activity^22^ and plasticity^21^ and might even be potentially filtered out by postsynaptic neurons. However, our observation that SS is selectively entrained by gamma rhythms suggests that SS effectively influences downstream neurons by precise millisecond gamma-band spike coordination across neurons^27,34,35,39^.

Place fields are a well-established property of CA1 neurons that support spatial navigation and memory. We found that CS firing rate carries more spatial information than SS^40,41^(but see^3^), and CS and SS code unique spatial information with CS place fields shifted backwards relative to animal’s movement direction. Furthermore, we found that CS strongly relates to intracellular theta power and dynamics^16^, suggesting CS is a critical substrate for phase precession^16,42^, another important spatial coding mechanism. In accordance, we also found that CS contributed stronger to theta phase precession coding relative to LFP theta than SS.

Our results provide direct experimental support that CS, and SB more generally, encode different circuit input features^38,43–51^. Differential roles of SS and SB in signal transmission have been found in the mammalian neocortex ^13,32^, thalamus^6,7,46^ and cerebellum^9,11,52^, the fly sensory system^49–51^ and electrosensory system of the electric fish^44,47,48^. The prevalence of SS and SB across species and structures suggest that these two distinct spike output codes are a general property of projecting neurons, which might have arisen from the need to enhance information transmission and computational power via multiplexing and parallel coding while constraining the number of neurons and axonal wiring^43–45,52–54^.

## Methods

All *animal experiments* were performed in accordance with the National Institute of Health Guide for Laboratory Animals and approved by the Boston University Institutional Animal Care and Use and Biosafety Committees.

### Mouse preparation

14 adult female C57BL/6 mice (Charles River Laboratories, Inc.), 8-12 weeks at the start of the study, were used for all experiments. Mouse preparation was as described previously^64– 66^.Custom recording apparatus consists an imaging window attached to an infusion cannula and a LFP electrode. The imaging window consists of a stainless steel cannula (OD: 3.17mm, ID: 2.36mm, 1.75mm height, AmazonSupply, B004TUE45E) with a circular coverslip (#0, OD: 3mm, Deckgläser Cover Glasses, Warner Instruments Inc., 64-0726 (CS-3R-0)) adhered to the bottom using a UV curable adhesive (Norland Products Inc., Norland Optical Adhesive 60, P/N 6001). We attached an infusion cannula (26G, PlasticsOne Inc., C135GS-4/SPC), and a stainless steel electrode steel wire for local field potential (LFP) recordings (Diameter: 130μm, PlasticsOne Inc., 005SW-30S, 7N003736501F) to the side of the imaging window using super glue (Henkel Corp., Loctite 414 and Loctite 713). The LFP electrode protruded from the bottom of the imaging window by 200μm, whereas the drug infusion cannula was level with the base of the imaging window.

Recording apparatus was surgically implanted under 1-3% isoflurane anesthesia. Analgesia was provided with sustained release buprenorphine hydrochloride (0.03 mg/kg, i.m.; Reckitt Benckiser Healthcare) administered preoperative that provides continued analgesia for 72 hours. A craniotomy of ∼3mm in diameter was made over the right dorsal CA1 (AP: -2mm, ML: +1.8mm). A small notch was made on the posterior edge of the craniotomy to accommodate the infusion cannula and the LFP recording electrode. The overlying cortex was gently aspirated using the corpus callosum as a landmark. The corpus callosum was then carefully thinned to expose the dorsal CA1. The imaging window was positioned in the craniotomy, and Kwik sil adhesive (World Precision Instruments LLC, KWIK-SIL) was applied around the edges of the imaging window to hold it in place. A small ground pin was inserted into the posterior part of the brain near the lambda suture as a ground reference for LFP recordings. Three small screws (J.I. Morris Co., F000CE094) were anchored into the skull, and dental cement was then gently applied to affix the imaging window, the ground pin, and an aluminum headbar posterior to the imaging window. See **Fig. 1a** for a diagram of recording apparatus placement.

AAV virus was infused via an infusion cannula (33G, PlasticsOne Inc., C315IS-4/SPC) connected to a microinfusion pump (World Precision Instruments LLC, UltraMicroPump3–4), through the implanted cannula. Infusion cannula terminated about 200um below the imaging window. 500 or 1000nL of AAVs were infused at a rate of 50-100nL/min, and then the infusion cannula was left in place for another 10 minutes to facilitate AAV spread. AAVs used were AAV9-Syn-SomArchon-GFP (titer: 5.9e^12^ genome copies (GC)/ml, UNC vector core), AAV9-Syn-SomArchon-GFP-p2A-CoChR (titer: 5.9e^12^ GC/ml, UNC vector core) and AAV9-Syn-SomArchon-BFP-p2A-CoChR (titer: 1.53e^13^ GC/ml, Vigene Biosciences, Inc).

### Rat preparation

4 male Long-Evans rats (Charles River Laboratories, Inc.), 350-500g, were used for all extracellular recording experiments. The surgical procedure was as described previously in detail^67^. Briefly, surgeries were performed under 1.5-3% isoflurane anesthesia (Webster Veterina Supply). Animals were injected with Buprenex (buprenorphine hydrochloride, 0.03 mg/kg, i.m.; Reckitt Benckiser Healthcare) and Cefazolin (330 mg/ml i.m.; West-Ward Pharmaceutical) preoperatively. Animals were implanted with custom unilateral microdrives containing 18-24 independently drivable tetrodes in the dorsal CA1 (AP: - 3.6mm, ML: +2.6mm, from bregma). Animals received postoperative Buprenex and Cefazolin two times a day for 3 days.

### Single neuron SomArchon voltage imaging

Habituated mice were head-fixed on a custom air-pressured spherical Styrofoam ball and free to run. Animals were recorded 3-4 weeks after surgery. All single cell SomArchon imaging was acquired via a customized widefield fluorescence microscope equipped with either a Hamamatsu ORCA Fusion Digital sCMOS camera (Hamamatsu Photonics K.K., C14440-20UP) for 828Hz voltage imaging, or an ultra-high-speed sCMOS camera (Kinetix, Teledyne) operating at 8bit mode for 5kHz and 10kHz voltage imaging. A 40x NA 0.8 water immersion objective (Nikon, CFI APO NIR) was used. A 140mW fiber-coupled 637 nm laser (Coherent Obis 637-140X) was coupled to a reverse 2x beam expander (ThorLabs Inc., GBE02-E) to obtain a small illumination area of ∼30-40 μm in diameter to minimize background fluorescence. A mechanical shutter (Newport corp., model 76995) was positioned in the laser path to control the timing of illumination via a NI DAQ board (USB-6259, National instruments). The laser beam was coupled through a 620/60nm excitation filter (Chroma technology corp.) and a 650nm dichroic mirror (Chroma technology corp.), and SomArchon fluorescence emission was filtered with a 706/95nm filter (Chroma technology corp.).

GFP or BFP fused with SomArchon was used for localization of SomArchon expressing cells during each recording. GFP visualization was with a 470 nm LED (ThorLabs Inc., M470L3), an excitation 470/25 nm filter, a 495 nm dichroic mirror and a 525/50 nm bandpass emission filter. BFP was visualized with a 395nm LED (ThorLabs Inc., M395L4), a 390/18nm excitation filter, a 416nm dichroic mirror and a 460/60nm emission filter. SomArchon fluorescence was either acquired at 828 Hz (16 bits, 2×2 binning), 5kHz or 10kHz, using HCImage Live (Hamamatsu Photonics). HC Image Live data were stored as DCAM image files (DCIMG) and analyzed offline with MATLAB (Mathworks Inc.). We did not detect notable differences for spike shape estimation for recordings performed at 5kHz versus 10kHz, and thus the imaging data were grouped together.

### Multi-neuron SomArchon voltage imaging with patterned illumination

To record multiple CA1 neurons, a custom digital micromirror device based targeted illumination microscope was used was described previously in detail^32^. In short, the custom-built widefield imaging scope includes a 6W 637 nm fiber-coupled multi-mode laser (Ushio America Inc., Necsel Red-HP-FC-63x), a digital micromirror device (DMD, Vialux, V-7000 VIS) and a high-speed and large sensor sCMOS (Hamamatsu, ORCA-Lightning C14120-20P). The laser output was collimated (Thorlabs, F950SMA-A), expanded (Thorlabs, BE02M-A), and directed onto the DMD. The DMD was controlled using a custom Matlab script based on Vialux ALP-4.2 API. SomArchon fluorescence was acquired at 500 Hz (12bit) with 1152 × 576 pixels (2×2 binning), corresponding to a 360 × 180 μm^2^ field of view. A subset of the CA1 multi-neuron imaging dataset was previously published ^32^ and was re-analyzed here.

### Optogenetics

To excite CoChR we used a blue 470 nm LED (ThorLabs Inc., M470L3) coupled to the laser path via a 416nm dichroic mirror, and controlled by T-Cube LED driver (ThorLabs Inc., LEDD18, lowest gain) modulated with MA TLAB (Mathworks Inc.) via a NI DAQ board (USB-6259, National instruments). A neutral density filter (ThorLabs Inc., ND13A, optical density 1) was used to reduce LED illumination density. Further, we used a pinhole (Olympus BX3-URA8 fluorescence illuminator turret) to reduce the illumination to a circular area of 55μm radius under the 40x objective. For SomArchon-GFP-p2A-CoChR recordings, we used a LED intensity range of 0.22-1.3mW/mm2. For SomArchon-BFP-p2A-CoChR recordings, we used a higher LED intensity range of 2.6-8.4mW/mm2, because neural CoChR response was less sensitive to blue light stimulation likely to lower CoChR expression. Simultaneous SomArchon and optogenetic recording trials consists of 1 second period before blue light illumination, 1.5 second blue light illumination period, followed by 0.5 second period without blue light illumination. The 1.5-second-long blue light illumination was either continuous, or pulsed at 40Hz (8.3ms per pulse) or 8Hz (8.3ms per pulse or 42ms per pulse). Inter-trial intervals were 5 seconds.

### Local field potential (LFP) recording

LFPs were recorded using the Open Ephys platform (http://open-ephys.org) at a 10 kHz sampling rate, filtered between 1Hz and 7.5kHz and then downsampled offline to 1kHz. To synchronize voltage imaging and LFP recordings during offline data analysis, Open Ephys system also recorded the TTL pulses that were sent by the sCOMS camera at the onset of voltage imaging data acquisition as well as a TTL pulse at the onset of each image frame. Additionally, to align optogenetic stimulation timing with recordings, we also recorded with Open Ephys System the voltage that was used to control the blue LED driver during optogenetics.

### Extracellular tetrode recording and animal tracking

Extracellular recordings from custom tetrodes were recorded by a OmniPlex D Neural Acquisition System (Plexon). Each channel was amplified and bandpass filtered (154 Hz to 8.8 kHz) to obtain both single-unit spike activity and LFPs (1.5 Hz to 400 Hz). Spike channels were locally referenced to remove both movement-related noise and potential electrical noise. Spikes were detected via threshold crossing and digitized at 40 kHz. To isolate single-units, waveform clusters from all four electrodes within a tetrode were manually identified using the Offline Sorter v3 (Plexon). Cineplex Studio (Plexon) was used for capturing animal location data via three infrared LEDs positioned atop the surgically implanted microdrive. Cineplex Editor (Plexon) was employed offline to enter event markers and to verify animal position data.

### Linear track environment

Rats were trained on a custom-built wooden linear track (223.5 cm long x 10.8 cm wide) elevated at 96.5 cm above the ground, as detailed previously^67^. Both ends of the linear track had water ports positioned. One end had a moveable wooden box with a sliding wooden door to restrict the access of the animal to the track during the 10sec long inter-trial-interval period. The linear track was situated in the middle of the room and partially enclosed, so that rats can use local and distal room cues to orient on the linear track.

### Behavioral training

Rats were first trained to run along the linear track to retrieve rewards at both ends of the track^67^. Once sufficiently trained, exhibiting reliable and consistent track running behavior, meaning rats left the home box when the door was opened and returned to home box to receive the second water reward. Rats were then implanted with recording device. After surgery and recovery, animals had refresher training sessions to assure reliable track running behavior. During behavioral experiments, the linear track length and the room lighting conditions were variable^67^. Here, we only analyzed data collected when rats ran on the longest track length with the room light on.

### Motion correction & neuron identification

SomArchon fluorescence images in DCMI format acquired by HCImage software were imported into Matlab. SomArchon fluorescence images were first motion corrected using a pairwise rigid motion correction algorithm as described previously ^68^. In short, the displacement of each image is computed by identifying the max cross-correlation coefficient between each image and the reference image. Our recordings consisted of multiple multi-second trials. For single-cell voltage imaging, each video file corresponding to one trial was first concatenated into a multi-trial image data matrix, and then we applied the motion correction algorithm. Since the illumination area is about 30-40μm in diameter, a rectangular window large enough to cover the entire neuron across all frames was selected manually for motion correction for single cell voltage imaging. The window selection was chosen to avoid large regions of dark parts of image and to include regions that had distinguishable contrasts that facilitate comparison with reference image. For multi-cell voltage imaging, due to image file size, we did not concatenate the data across trials. Each trial was first motion corrected individually. We then corrected motion offsets across trials by referencing all trials to the first trial. The motion-corrected image sequences were then used for subsequent manual ROI neuron identification using the drawPolygon function in Matlab. SomArchon fluorescence traces for each ROI were then extracted from the motion-corrected image sequences. Background fluorescence was estimated by averaging all pixels that are not part of the ROI. The traces were then detrended to correct for photobleaching by subtracting a low-pass filtered version of the trace (1.5sec rectangular smoothing kernel). The resulting fluorescence traces were then used for subsequent analysis.

### Spike detection, subthreshold Vm trace extraction and spike signal-to-baseline ratio (SBR) calculation

Spike detection was performed similar to that described previously in Xiao et al^32^. To estimate baseline, we first estimated baseline fluctuations by averaging the fluorescence trace using a moving window of ±100 frames to obtain the “Smoothed Trace” (ST). We then removed potential spike contributions to the baseline line by replacing fluorescence values above ST with the corresponding values of ST resulting in a spike-removed trace including only the subthreshold baseline fluctuation. To identify spikes, SomArchon fluorescence traces were high-pass filtered (>120Hz), and then spikes were detected as a fluorescence change greater than 4 standard deviations of the baseline subthreshold fluctuations.

To extract subthreshold membrane voltage (Vm) fluctuations, we removed three data points centered at the peak of each detected spike from non-filtered SomArchon dF/F trace and interpolated the missing data points with the surrounding data points. To calculate spike signal-to-baseline ratio (SBR), we first obtained the spike amplitude by calculating the difference between the peak spike fluorescence and the lowest fluorescence value within three data points prior to the spike. We then divided the spike amplitude by the standard deviation of the Vm across the entire recording duration. To assure signal quality, only neurons with an averaged SBR of at least 4 were included for analysis.

### Complex spike (CS) and single spike (SS) detection

A common way to identify bursting in extracellularly recorded single-unit data is to identify a cluster of spikes with short inter-spike intervals (ISIs). However, ISI distributions for individual neurons are often continuous, and thus it is difficult to determine a definitive ISI threshold, leading to variation in ISI threshold of ∼6-14ms across studies. Recent intracellular studies demonstrated that individual spikes with CS can have ISIs of more than 10ms^20^. Thus, we used both ISI and Vm after-depolarization potential (ADP) to classify CS and SS. CS were detected as follows: First, putative spike bursts were detected based on ISI 14ms criterion and we classified each spike within a putative burst in terms of their order (1^st^ spike, 2^nd^ spike, …). ADP was calculated by the difference in the mean Vm during the 5-25ms period after the 1^st^ burst spike and the mean Vm during the 5-25ms before the 1^st^ burst spike, normalized by the averaged spike amplitude across the entire recording duration. If a putative burst had an ADP of more than 15%, it was considered as CS and all spikes of the burst was classified as CS spikes. Spikes that are not identified as CS were considered SS. SS could hence have ISI smaller than 14ms but are conceptualized here as high firing rate but regular spiking events.

### Spectral decomposition

Spectral decomposition of SomArchon Vm or LFP was performed with FieldTrip Matlab toolbox^69^ (https://www.fieldtriptoolbox.org/), using wavelet morlets functions (5 cycles). The complex wavelets coefficients, from which we extracted the phase or power, were used to compute spike-LFP/Vm phase locking values, and to estimate of the spectral power.

### Phase locking strength (PLS)

To obtain a quantification of how consistent spikes occur relative to the phase of an oscillation we calculated the phase-locking value (PLV^70^) defined as:

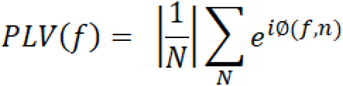

where f is frequency and N being the total number of spikes. The phase ϕ was obtained from the complex wavelet spectrum.

Since PLV is not independent of the number of spikes considered and tends to inflate with low spike numbers. We only included neurons that had at least 10 spikes for spike-PLV analysis. Further, we adjusted the PLV value using the following equation^71,72^ (mathematically equivalent to pairwise phase consistency) to account for any potential difference in the number of CS and SS detected in a neuron, which we term here as phase locking strength (PLS):

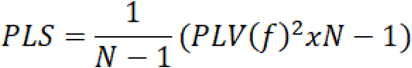

where N is the number of spike occurrences and f is frequency.

Finally, to further reduce any residual spike influence on PLV estimates from spike-removed SomArchon Vm traces, we shifted the spike train and the corresponding Vm trace by 8.2ms (7 frames at 828Hz sampling rate) to each other, which reduced the PLS inflation in the higher frequency range (>100Hz) due to spike number.

### Extracellular tetrode recordings

#### Spike burst (SB) detection

Given that we did not have access to membrane potentials for extracellular tetrode recordings, we identify SBs as periods of high frequency spiking with ISI less than 10ms. We chose a more conservative (smaller ISI) criterion for identifying SB than for identifying CS from voltage imaging recordings, because we do not have ADP-based criterion. Spikes not considered being part of any SB were defined as single spikes. We only included neurons with mean firing rates below 20Hz estimated over the whole recording, because there was distinct subset of neurons, presumably fast-spiking interneurons, that exhibited very high tonic firing rates (small ISI) and thereby difficult to classify with a ISI-based criterion alone.

#### CA1 place fields

We followed an approach as proposed by Skaggs^7^. We did not attempt to separate the CA1 neural population into place cells and non-place cells, but we characterized the amount of spatial information using information theoretical measures for each neuron and as a function of spike bursts and single spikes.

#### Spatial information and sparsity index

The amount of information a neuron had about the animal’s position was defined as^7,73^:

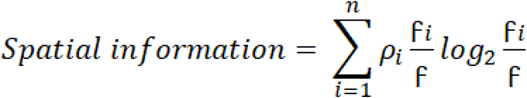

Where p_i_ is the position bin occupancy defined as p_i_= t_i_/∑t_i_ (summed over i) and where f is the mean firing rate defined as f=∑p_i_f_i_.

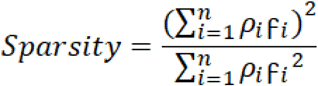

Where f_i_ is firing rate of the neuron at the ith position bin.

#### Center of mass (COM) calculation

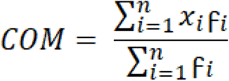

where x_i_ is the ith position bin on the linear track and f_i_ is firing rate of the neuron at the ith position bin.

#### Quantifying the relationship between Spike-LFP theta phase and position on linear track

For the place field theta spike phase analysis, we filtered the LFP signal in the theta (3-12Hz) frequency range using a butterworth filter kernel. To obtain the instantaneous phase, we used the analytical signal of the Hilbert transform. To quantify the relationship between a neuron’s preferred spike-LFP theta phase and animal’s position, we applied circular-linear correlation^74^ that estimates a coefficient of multiple correlation (analogues to the coefficient of determination, which is the square of the correlation value) ranging from 0 to 1 with 1 meaning all the variance is shared and 0 meaning no variation is shared.

## Resource availability

## Lead Contact

Further information and requests for code and data should be directed to the lead contact Xue Han or Eric Lowet.

## Materials availability

This study did not generate new unique reagents.

## Data and code availability

- Data are available from lead contact upon request.
- Codes used for data analysis is available on Github repository: https://github.com/HanLabBU.
- Any additional information required to reanalyze the data reported in this paper is available from the lead contact upon request.

## Extended data figures

**Extended data Fig.1:**
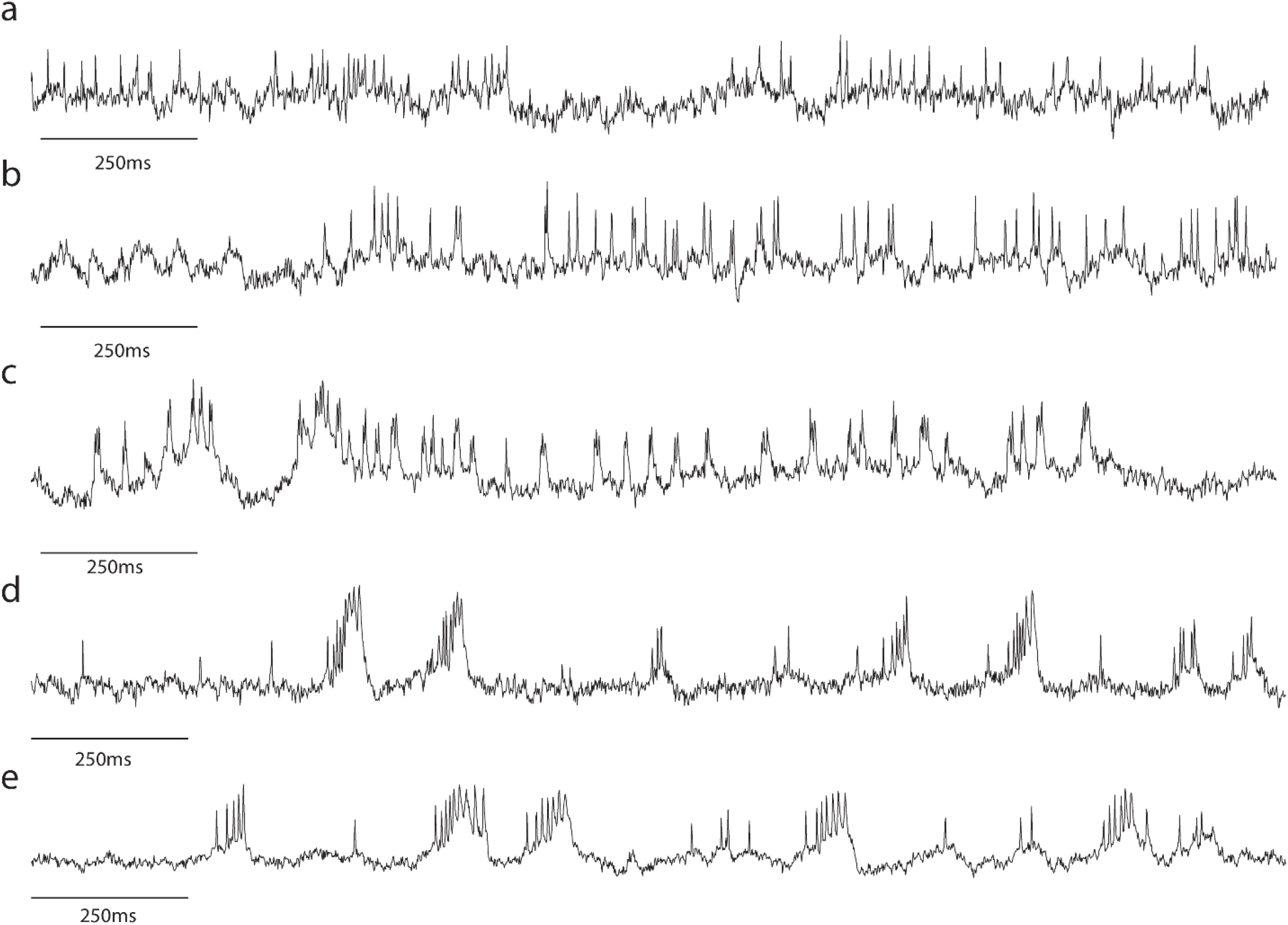
CA1 SomArchon membrane voltage trace examples. (**a**). A neuron with no spike bursts (**b**). A neuron with few short bursts (**c**). A neuron with frequent short bursts and membrane depolarization (**d**). A neuron with large bursts and steep membrane depolarization. (**e**). A neuron with particularly long bursts and large membrane depolarization.

**Extended data Fig.2:**
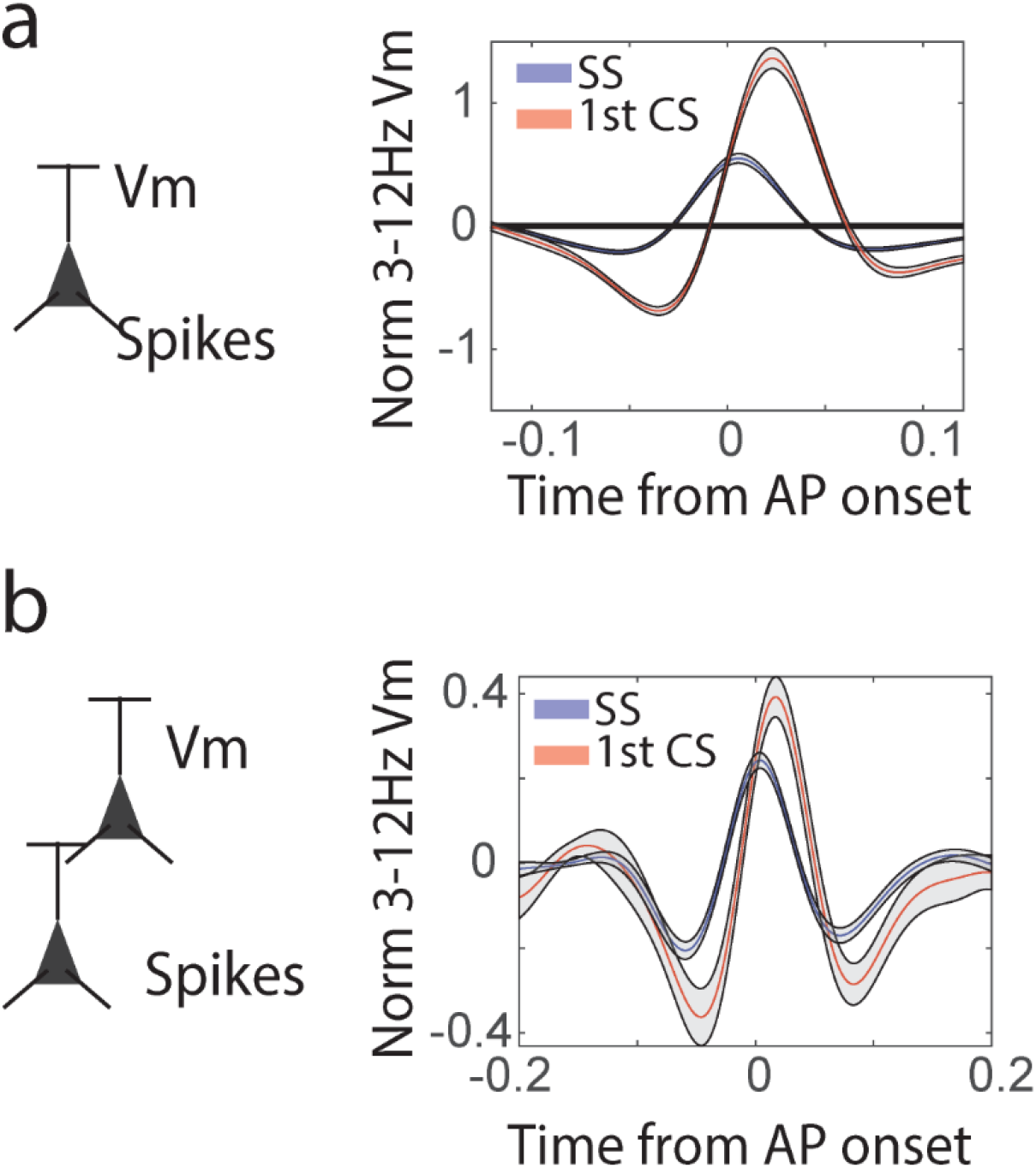
Theta frequency (3-12Hz) filtered Vm aligned to spikes of individual neurons recorded under different imaging conditions. **(a)** Single neuron SomArchon imaging condition. Theta frequency (3-12Hz) filtered Vm aligned to spikes for either SS (blue) or CS (red). The first CS spike was used as reference spike. The spike-triggered windows were then averaged. **(b)**. Same as a, but for multiple neuron SomArchon imaging condition. Only neurons that were at least 40microns apart were included in this analysis.

**Extended data Fig.3:**
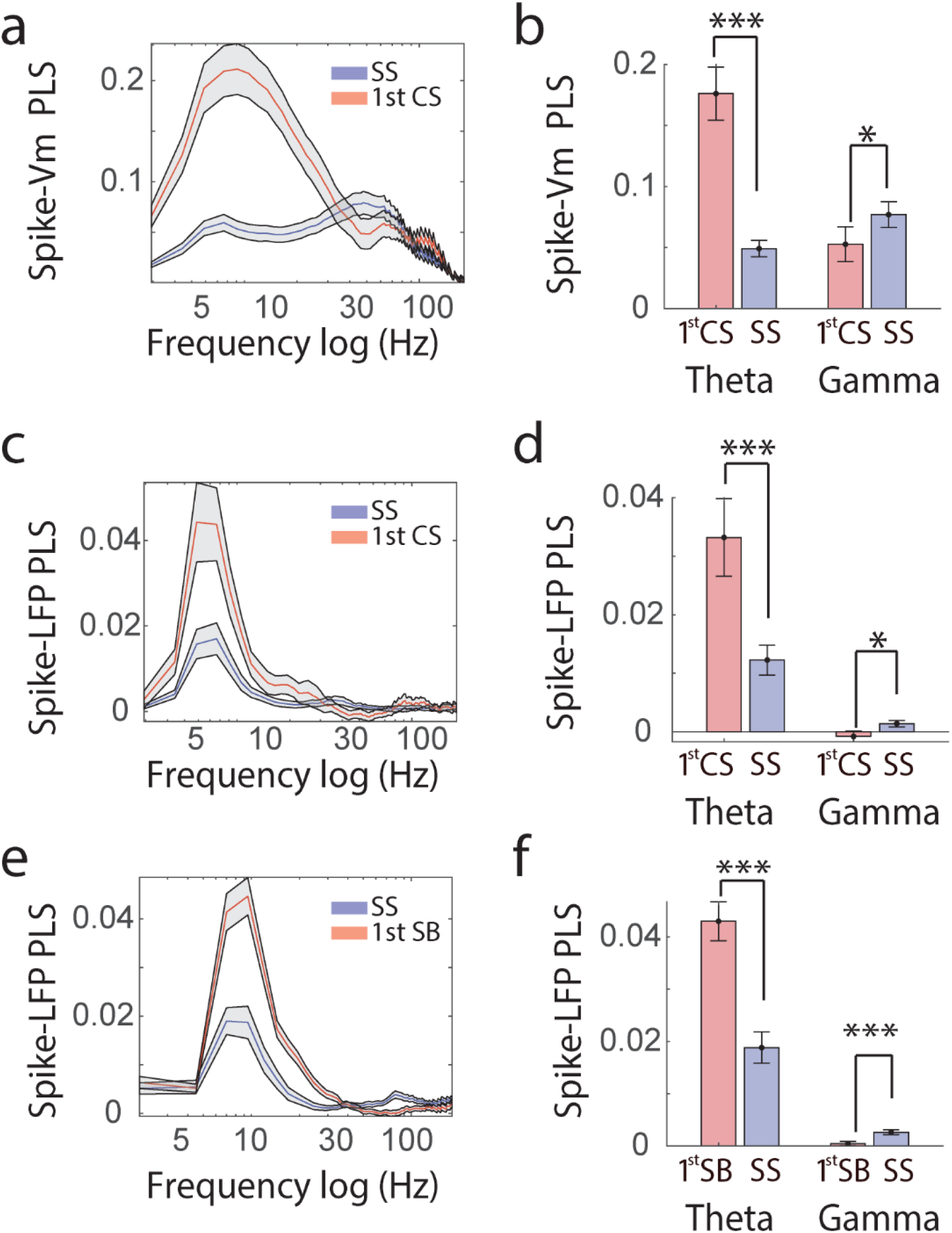
Comparison of spike PLV to Vm and LFP between SS and 1^st^ CS recorded in mouse CA1 (a-d) and 1^st^ SB recorded in rat CA1 (e-f). **(a)**. PLS of spikes to Vm across frequencies for the population of recorded neurons (**b)**. PLS of spikes relative to Vm theta (3-12Hz) and gamma oscillations (30-90Hz) for CS (red) and SS (blue). Spike-Vm theta PLS for CS is greater than for SS (paired t-test, p=5.8e^-9^, df=37). Spike-Vm gamma PLS for SS is greater than CS (paired t-test, p=0.017, df=37). **(c)**. PLS of spikes to LFP across frequencies for the population of recorded neurons (**d)**. PLS of spikes relative to Vm theta (3-12Hz) and gamma oscillations (30-90Hz) for CS (red) and SS (blue). Spike-Vm theta PLS for CS is greater than for SS (paired t-test, p=1.86e^-4^, df=37). Spike-Vm gamma PLS for SS is greater than CS (paired t-test, p=0.01, df=37). **(e)**. PLS of spikes to LFP across frequencies for the population of recorded neurons (**f)**. PLS of spikes relative to Vm theta (5-12Hz) and gamma oscillations (40-100Hz) for CS (red) and SS (blue). Spike-Vm theta PLS for CS is greater than for SS (paired t-test, p= 5.41e^-13^, df=164). Spike-Vm gamma PLS for SS is greater than CS (paired t-test, p=1.6e^-4^, df=164).

**Extended data Fig.4:**
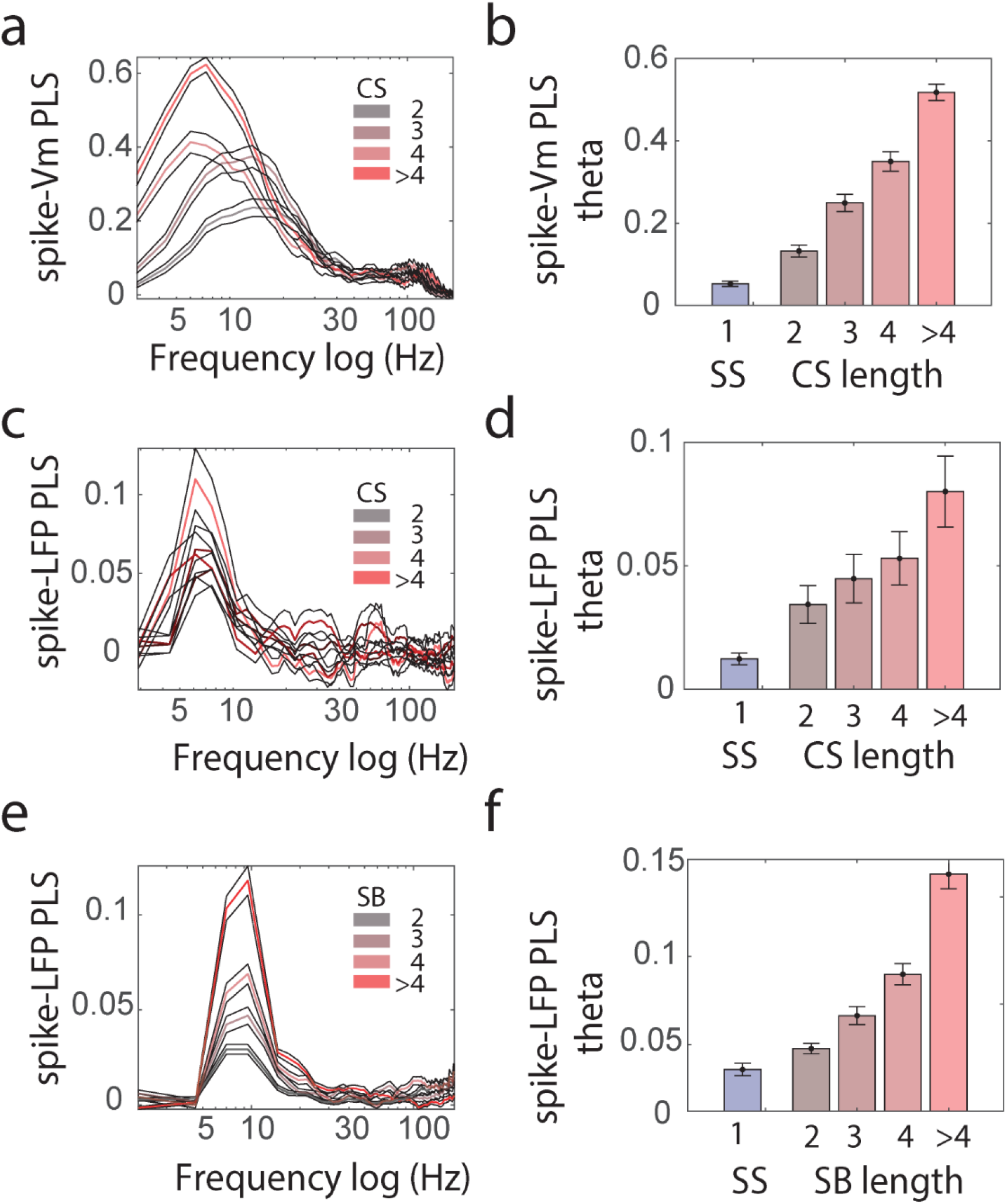
Effect of burst length on theta phase locking. **(a-d)** Mouse CA1 recordings. **(e-f)** Rat CA1 recordings. **(a)**. Spike-Vm PLS for CS containing 2, 3, 4 or >4spikes (colored from darker to lighter color). **(b)**. Quantification of Spike-Vm theta PLS for SS and CS containing increasing spike numbers shown in **a**. Spike-Vm theta PLS is increase for CS with more spikes (linear regression, r^2^=0.36, p=1e^-20^, n=156). **(c)**. Spike-LFP PLS for CS containing 2, 3, 4 or >4spikes (colored from darker to lighter color). **(d)**. Spike-LFP theta PLS is significantly higher for CS with more spikes (linear regression, r^2^=0.36, p=0.0369, n=156). **(e)**. PLS of SB relative to LFP across frequencies versus the number of spikes within a SB. Blue is SS. Dark to lighter red colors reflect SB containing 2, 3 4, and >4 spikes. **(f)**. Quantification of PLS relative to LFP theta frequencies for SB with varying number of spikes (linear regression slope, p<1e^-20^, n=66).

**Extended data Fig.5:**
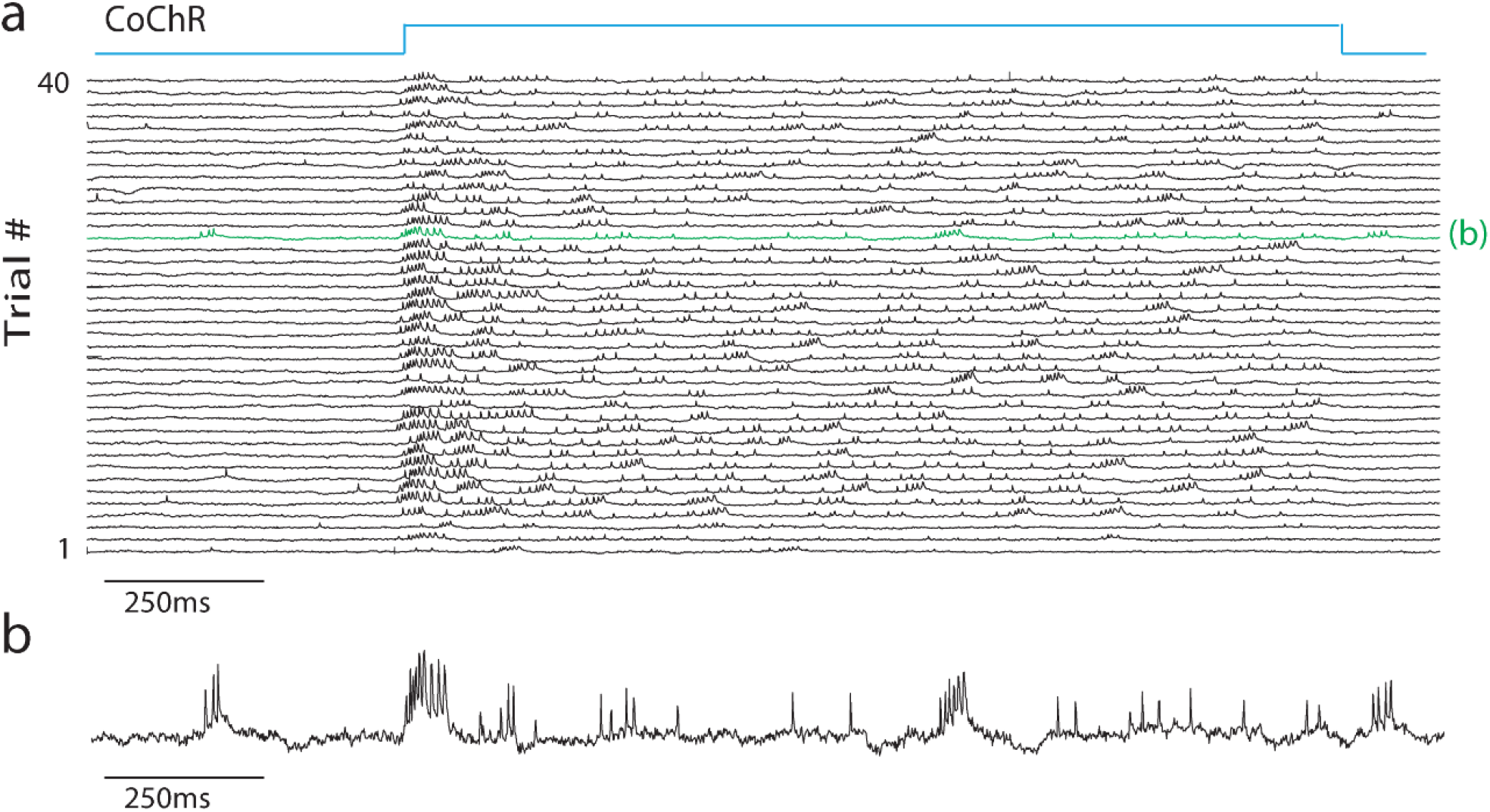
An example CA1 neuron recording showing SomArchon fluorescence traces upon 1.5 second continuous optogenetic stimulation of CoChR. **(a)**. SomArchon fluorescence traces across 40 trials. Optogenetic stimulation period is illustrated by the blue light on the top. In this particular neuron, there was an increase in SS and CS spike rate upon CoChR stimulation. (**b**). Zoom in view of an example trial, the trial colored in green in **a**.

**Extended data Fig.6:**
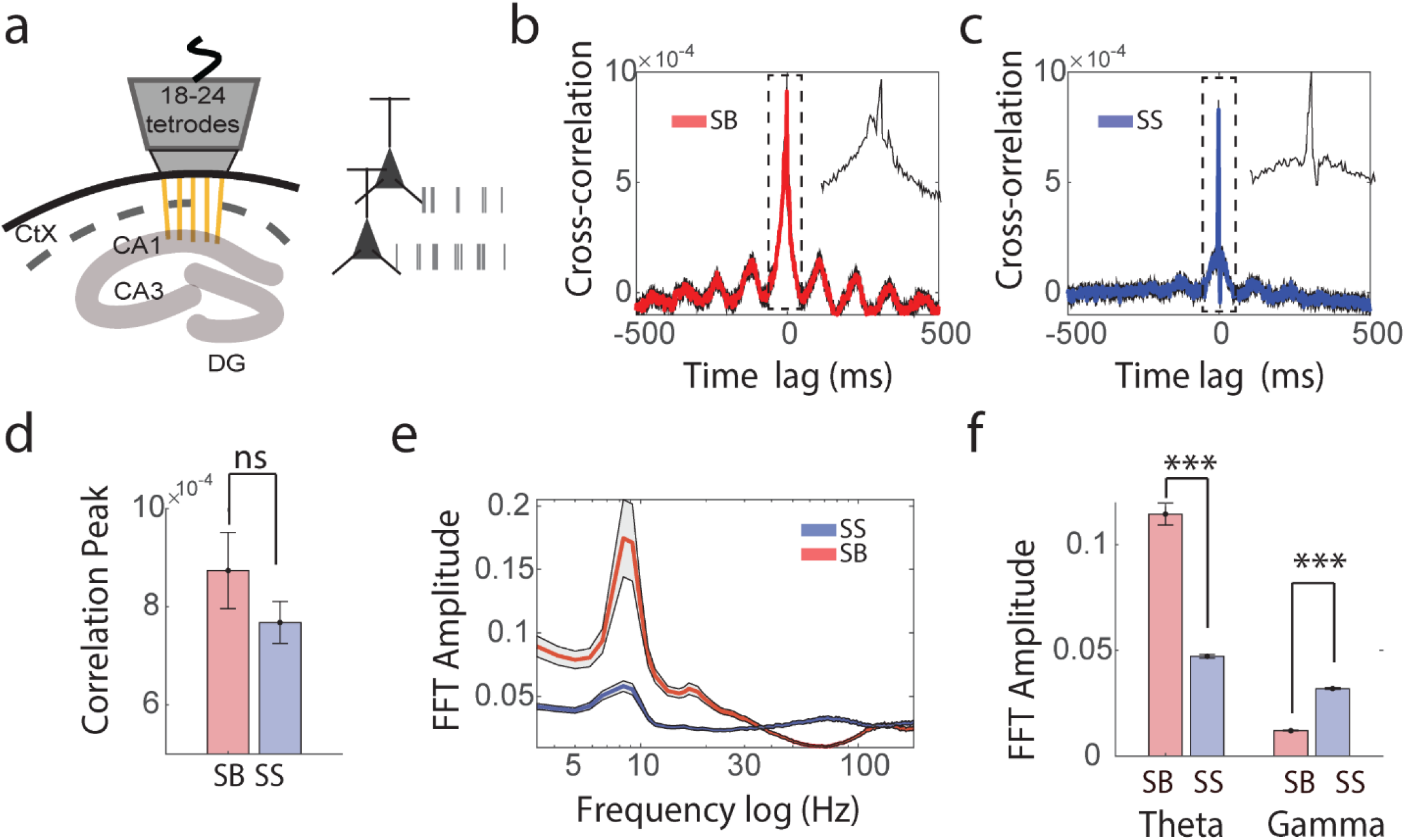
Power spectrum component of spike train cross-correlograms for SS and SB recorded in rats. **(a)**. Schematic representation of the extracellular recording setup containing between 18-24 tetrodes in the rat dorsal CA1. **(b-c)**. The mean population averaged spike cross-correlogram for SB (**b**, red) and SS (**c**, blue) between simultaneously recorded neuron pairs (n=1608). Inset is a zoom-in on the central peak of the cross-correlogram (−50 to +50ms) **(d)**. Quantification of the spike cross-correlation peak between SB and SS (paired t-test, p=0.1, df=1607). **(e)**. The FFT amplitude spectrum of the spike cross-correlograms (red=SB, blue=SS). **(f)**. Quantification of the FFT amplitude of the cross-correlogram in the theta-band (paired t-test, p = <1.e^-20^, df=1607) and gamma-band (paired t-test, p = <1.e^-20^, df=1607).

